# Filling up the tree: considering the self-organization of avian roosting behavior

**DOI:** 10.1101/000349

**Authors:** Bradly Alicea

## Abstract

In this paper, models for understanding bird roosting will be considered for purposes of developing better Artificial Life models of complex behavior. Roosting involves multiple flocks of birds picking a single tree limb to rest on for the night, and can be considered an iterative, time-dependent process that unfolds over a 45-minute interval roughly corresponding to twilight. Two models will be used to better understand the main components of this behavior. The constrained dynamics model, which represents continuous random absorption on a one-dimensional lattice, will be used to characterize the dynamics of crowding in the tree structure over time. A second approach involves the relationships between complex networks and roosting behaviors, in particular the evolution of structured networks via rules of incorporation and interaction. Finally, the percolation model will be proposed as a way to bridge behaviors explained by the constrained dynamics and complex network models.

## Introduction

While there are a number of ways to model and simulate collective animal behavior, many of these models are related to the emergent properties of flocking and swarming behaviors. Other behaviors (notably bird roosting) can be used to better understand the full range of collective animal behaviors as well as develop better simulation of the interactions between an animal’s biological substrate (e.g. brain) and behavior. Implementations of these so-called artificial life models use both formal data structures and statistical models. Here, two descriptive models and one simulation-based model will be introduced. The constrained dynamics and network growth models will define specific aspects of roosting behavior. The percolation model will reproduce each of these behavioral aspects in a way tractable to simulation.

Paradigms that have attempted to simulate animal behavior have typically used simple interaction rules to produce emergent phenomena. One early example called “boids” simulated bird flocking behaviors (Reynolds, 1987). The “boid” system used several collision detection parameters to produce flocking behaviors. What resulted was a system that produced an abundance of intricate patterns called weak emergence, but a paucity of intelligent, self-directed collective behaviors. Later attempts included swarm intelligence approaches, which relied upon physical links between agents interacting in the system. For example, Gambardella and Dorigo (1995) were able to solve an NP-hard optimization problem using a network of virtual pheromone trails and a population of virtual ants who followed only the most densely concentrated paths. In this paper, a complex, coordinated behavior will be modeled using more complex computational methods than have previously been employed. These insights will then be used to potentially inform the development of a novel artificial life system.

Roosting among Florida Egrets (*Egreta thula*) is a complex behavior that involves multiple flocks of birds picking a single tree limb to rest on for the night. The process unfolds as follows: there is initial placement by the first incoming wave of birds, followed by interactions between the roosted group of birds and a subsequent wave of immigrants. These interactions result in the relocation of ‘displaced’ members from the first wave. This process of immigrant groups arriving at the tree and continually displacing established roosters continues until all waves of incoming birds have been assimilated, all available branches are taken, or the roosting period ends. Thus, roosting can also be considered a *time-dependent process* in that it unfolds over a 45-minute interval that roughly corresponds with twilight. The rate at which incoming waves occur and the total number of birds in all flocks act as constraints.

## “Nodes” of the Tree

This section of the paper will discuss the best way to characterize interactions between elements in this system. A review of relevant natural history will inform a formal strategy for modeling the roosting surface.

### Natural History of Roosting

According to Beauchamp (1999), there are three main benefits to participating in roosting: a reduction in thermoregulative demands on the individual, a decrease in predation risk for the individual, and an increase in foraging efficiency for certain members of the population. Generally speaking, the ecological causes of roosting are mostly unknown. However, in cases where such communal behaviors are costly to maintain and perform, changes in the local environment can lead to reduced aggregation and may induce solitary behavior (Wcislo and Danforth, 1997).

Roosting has been explained in the past by the “information center” hypothesis (see Ward and Zahavi, 1973). Typically, this explanation has been contingent upon two parameters: foraging success and social rank. For example, not all members of the flock are equally successful at foraging. Less successful conspecifics can recognize successful foragers, and will inadvertently follow them from food patches to roosting sites. In this way, the incoming waves of egrets may represent miniature dominance hierarchies that readjust themselves as new birds integrate themselves into the system.

From a behavioral ecology standpoint, it has been observed that roosting birds form a structured assemblage once a given population is entirely settled onto a roosting surface (Weatherhead, 1983). Hamilton (1971) has observed that flocks of birds position themselves optimally with regard to social rank. For example, dominant birds will tend to centralize themselves spatially and surrounded themselves with subordinates (deGroot, 1980).

### Modeling the Roosting Surface

A basic problem in terms of understanding how birds roost in a tree over a finite time period is how global processes play a role in local behavior and interactions. One way of thinking about bird immigration as a set of roosts is in terms of particles gradually populating a discrete surface. A common mathematical model for the problem of detecting general patterns of settlement from waves of migration is the *constrained dynamics model* (Kolan *et al*, 1999). This model represents random absorption that is continuous in time on an *n*-dimensional lattice (Jin *et al*, 1994). In the case of the roosting problem, a set of identical particles (birds) can absorb onto a surface (branch tips with discrete distributions in space) at a specific rate (*m*) per unit of surface area. Birds can also leave the surface at a rate of *n*, but the size of this parameter is usually infinitesimal (Evans, 1993). In addition, adsorption is subject to free volume constraints so that, as in a naturalistic setting, only one bird can occupy a single branch at a specific point in time.

In terms of behavioral dynamics (e.g. interactions between individuals), this can act as a source of scarcity and point of contention in the system. This could be related to energetic and other constraints. However, it may also be that low levels of density encourage a spreading out across the surface, so that new arrivals result in the migration of established individuals to empty roosts without any contention. Regardless of individual behaviors, a fundamental question remains: what is the critical degree of density (globally) at which individuals switch from being amicable to largely agonistic? Part of the answer may lie in the findings of Renyi (1958), who calculated that such a system “jams” when the density (*pc*) reaches .75. In general, a constrained dynamics-type system approaches a ‘jammed’ state according to the following formula

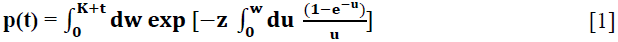

This model [1] also exhibits two distinct time scales at high values of *K* where *K* = *m*/*n*. Thus, high values of *K*, especially at later moments in the process, will result in high densities of birds in the tree. This can be used to explain what has been casually observed in nature; namely, that the decision to either share roosts with newcomers or fight is governed by a subtle process of global crowding. With the constrained dynamics model, we can only uncover general trends in the roost structure. However, the constrained dynamics model also provides us with a basis for further modeling of the process. Once we understand the various temporal and spatial dependencies of immigration process and the tree surface, then we can consider strategies and interactions engaged in by individual birds.

## “Arcs” of the Tree

This section of the paper will discuss the best way to characterize the general movement dynamics for elements in this system. A review of strategies employed by individual birds will lead to consideration of complex network dynamics that provide complex dynamics to this behavior.

### Strategy Deployment

All birds can be observed in nature to exhibit a fixed set of behavioral strategies when roosting. These were originally defined as “warding off”, “capitulating”, and “beak fencing”. If we assume that birds follow dominance hierarchy cues during the roosting process, the probability distribution of strategies will be skewed so that certain behaviors are more probable under particular conditions. For example, it was found from the behavioral data that “beak-fencing” was not a dominant strategy, suggesting that when space exists in the tree as predicted by the constrained dynamics model, the birds are willing to distribute themselves by “non-violent” means. However, beak fencing can occur relatively early on in the roosting process.

This result suggests, as would be predicted with the constrained dynamics model, that birds distribute themselves asymmetrically about the tree. They may even go as far as fighting to roost in certain areas, although the relative infrequency of beak fencing suggests that either many of these “prime” roosting areas exist, or that the constrained dynamics model is only partially predictive and thus only a few “prime” roosting spots actually exist. To verify which prediction is correct, we can use a complex graph to model the consequences of interactions between birds.

### Birds on a Wire (Complex Graphs)

Complex networks have been used to model everything from the migration patterns of dollar bills (Brockmann *et al*, 2006) to the parallel function of metabolic pathways. The edges of a graph based on roosting behavior are determined by sets of interactions between pairs of birds. Roosting graphs can also be considered complex networks, as bird migration within a tree ultimately results in a discontinuous graph with a potential for small-world clustering (see Watts and Strogatz, 1998). The strategies defined in the last section serves to give each bird a score, and when two birds “interact”, they both influence each other’s score and create a link between their respective branches. In this way, each link in the network represents a “flow” across the tree. As birds move around within the tree, they extend this graph even further, connecting all the perches they have previously visited. They also create a “degree” of connectedness; each bird can be directly connected to a number of other birds, and indirectly connected to infinitely more birds. However, it is expected that some birds will interact more than others.

As a result of this asymmetry, various degrees of connectivity will develop over time. Network “growth” is then generally defined by either incoming birds or newly interacting birds already in the tree. As the roosting process continues and the roosting surface becomes progressively filled, the network can be expected to grow at a faster rate. Given these internal migration paths and scores, we can model all of the interactions across the tree and the entire roosting session. Specifically, by combining the number of edges converging upon commonly contested perches and comparing the scores of the birds that occupy those perches, we can detect “hotspots” or “hubs” within the network topology that are particularly diagnostic of a small-world network.

Like all complex network models, we need a simple set of conditions governing the incorporation of newcomers to the network. Assuming that the roosting network is scale-free, or no characteristic number of connections per node, there are many potential attachment rules in our roosting network, and most likely several of them operate simultaneously. Attachment rules, such as the preferential rule of the Barabasi-Albert model (Newman, 2003; Barabasi *et al*, 2000), serve to selectively constrain a network as it grows and imposes spatial structure on collective behavior (Bornholdt and Schuster, 2003). The first of these is a dominance hierarchy within the community of birds, and it based on the Barabasi-Albert model. The preferrential attachment aspect of the Barabasi-Albert model may be used to understand the relationship between the spatial distribution of birds in the tree and their behavioral strategies. A second potential attachment rule is that of iterative observation. This can be made into a process using the “copying model” of Bollobas *et al* (2001).

In this scenario, incoming birds observe where the previous migrants have attempted to settle, and then mimic this behavior. However, subsequent movement within the tree structure and strategies employed during interaction are not copied. This may explain the relative lack of fighting for certain roosts, and the equal number of observations for warding off by the perched bird and capitulation by the perched bird. A third potential rule may involve a quasi-stochastic criterion for selecting places to roost. The Cooper-Frieze model (see Bornholdt and Schuster, 2003) might characterize this scenario; this algorithm combines features of preferential and uniform attachment. This may also explain the relative lack of crowding in certain portions of the tree; birds may just select places that have relatively few neighbors. Ultimately, we are interested in observing phase transitions within the graph structure that occur during the evolution of complexity.

## Toward a synthetic model of brain and behavior: what we have learned

The models presented here provide the basis for understanding complex behavior in a rigorous manner. However, we can consolidate the modeling of both initial arrival patterns and the interactive components of roosting in a percolation model (see Grassberger and Kantz, 1991). Percolation is a geometric process, and is not strictly concerned with fitness or energetics (Adami, 1998). While this may seem to be problematic given the natural history of roosting behavior, such a model holds everything else constant and allows us to focus on the landscape upon which this roosting behavior is playing itself out. Furthermore, a percolation model provides an easy way to investigate both the “connectedness” of a given space and its history. This allows for the simulation of network growth models as they relate to the spatial dynamics of roosting.

Percolation models consist of a finite lattice, the elements of which are selectively connected together via bonds. Whether or not the roosting model is extensible to other collective behaviors may be debated; however, the model proposed here is predicted to give a more realistic account of such behavior (Keitt and Johnson, 1995). The main evidence for this comes from the behavior of percolation models as they approach an infinite cluster. An infinite cluster connects all edges of the topology, and is determined by bonds between points in the lattice surface that exist with a probability of *p*. An infinite cluster exists once the strength of all bonds between lattice elements exceeds this probabilistic threshold. As this state is approached, the properties of behavior in a space can change drastically. This is what is called a geometric phase transition. Phase transitions in percolation models are similar to phase transition behavior exhibited by complex graphs as they grow beyond a certain size. According to Erdos and Renyi (see Bornholdt and Schuster, 2003), complex graphs undergo a phase transition around *t* ∼ *n/2* during complex graph evolution. This has been referred to as the emergence of a giant component, and may be related to the onset of jamming as characterized by the constrained dynamics model.

## References

Adami, C. (1998). Introduction to Artificial Life. Berlin, Springer.

Barabasi, A-L., Albert, R., and Jeong, H. (2000). Scale-free characteristics of random networks: the topology of the world-wide web. Physica A, 281, 69–77.

Beauchamp, G. (1999). The Evolution of Communal Roosting in Birds: origin and secondary losses. Behavioral Ecology, 10(6), 675–687.

Bollobas, B., Riordan, O., Spencer, J., and Tusnady, G. (2001). The degree sequence of a scale-free random graph process. Networks and Algorithms, 18(3), 279–290.

Bornholdt, S. and Schuster, H.G. (2003). Handbook of Graphs and Networks: from the genome to the internet. Wiley-VCH, Weinheim, Germany.

Brockmann, D., Hufnagel, L., and Geisel, T. (2006). The scaling laws of human travel. Nature, 439, 462–465.

deGroot, P. (1980). Information Transfer in a Socially Roosting Weaver Bird (*Quelea quelea*: Ploceinae): an experimental study. Animal Behavior, 28, 1249–1254.

Evans, J.W. (1993). Random and cooperative sequential adsorption. Reviews in Modern Physics, 65, 1281.

Gambardella, L.M. and Dorigo, M. (1997). Ant Colonies for the Traveling Salesman Problem. *IEEE* Transactions on Evolutionary Computation, 1(1), 52–66.

Grassberger, P. and Kantz, H. (1991). On a forest fire model with suppressed self-organized criticality. Journal of Statistical Physics, 63, 685.

Hamilton, W.D. (1971). Geometry of the selfish herd. Journal of Theoretical Biology, 31, 295–311.

Keitt, T.H. and Johnson, A.R. (1995). Spatial heterogeneity anomalous kinetics: emergent patterns in diffusion-limited predator-prey interaction. Journal of Theoretical Biology, 172, 127–139.

Kolan, A.J., Nowak, E.R., and Tkachenko, A.V. (1999). Glassy behavior of the constrained dynamics model. Physical Review E, 59(3), 3094–3099.

Jin, X., Tarjus, G., and Talbot, J. (1994). An adsorption-desorption process on a line: kinetics of the approach to closest packing. Journal of Physics A, 27, L195.

Newman, M.E.J. (2003). The structure and function of complex networks. SIAM Review, 45(2), 167–256.

Rényi, A. (1958). On a One-Dimensional Problem Concerning Random Space-Filling. Publications of the Mathematical Institute of the Hungarian Academy of Sciences, 3, 109–127.

Reynolds, C. (1987). Flocks, Herds, and Schools: a distributed behavioral model. Computer Graphics 21(4): 25–34.

Ward, P. and Zahavi, A. (1973). The Importance of Certain Assemblages of Birds as “Information Centers” for Food Finding. Ibis, 115, 517–528.

Watts, D.J., and Strogatz, S.H. (1998). Collective dynamics of ‘small-world’ networks. Nature, 393, 440–442.

Wcislo, W.T. and Danforth, B.N. (1997). Secondary Solitary: the evolutionary loss of social behavior. Trends in Ecology and Evolution, 12, 468–474.

Weatherhead, P.J. (1983). Two Principle Strategies in Avian Communal Roosts. American Naturalist, 121(2), 237–243.

